# Robustness of field studies evaluating biodiversity responses to invasive species management in New Zealand

**DOI:** 10.1101/2022.03.10.483864

**Authors:** Robert B. Allen, David M. Forsyth, Darryl I. MacKenzie, Duane A. Peltzer

**Affiliations:** Independent Researcher, 8 Roblyn Place, Lincoln 7608, New Zealand; Vertebrate Pest Research Unit, NSW Department of Primary Industries, 1447 Forest Road, Orange, NSW 2800, Australia; Proteus, Outram 9019, New Zealand; Manaaki Whenua Landcare Research, Lincoln 7608, New Zealand

**Keywords:** evidence-base, experimental design, logistic regression, pest control, randomisation, replication, representativeness, sampling universe, systematic review

## Abstract

Benefits of invasive species management for terrestrial biodiversity are widely expected and promoted in New Zealand. Evidence for this is presented in policy and scientific reviews of the literature, but the robustness and repeatability of the underpinning evidence-base remains poorly understood. We evaluated the design of field-based studies assessing biodiversity responses to invasive species management in 155 peer-reviewed articles published across 46 journals from 2010 - 2019. Each study was assessed against nine principles of experimental design, covering robustness of sampling and avoidance of bias. These principles are important in New Zealand to detect treatment effects from environmental variability driven by underlying gradients such as soil fertility, climate and disturbance. Fifty two percent of studies defined a sampling universe and 68% of studies specified the treatment. Whereas, 54%, 74%, and 50% of studies did not utilise replication, representatively sample the universe, or quantify invasive species, respectively. Ninety five percent of studies quantified biodiversity responses, although a high proportion of these did not representatively sample replicates. Initial conditions and accounting for effects of experimental implementation were not utilised in 57% and 84% of studies respectively. No studies avoided observer/analyst bias using blinding methods, despite this being widely adopted in other fields. Ordinal logistic regression showed these principles varied in how robustly they were applied among categories of biodiversity responses and invasive species. Our findings suggest that greater attention to experimental design principles is desirable: supported by researchers, funding agencies, reviewers, and journal editors. Greater resources is not necessarily a solution to these design issues. Undertaking fewer studies, that are individually more expensive because they better adhere to experimental design principles, is one alternative. Our intent in this article is to improve the robustness of future field studies for at least some principles. Robust designs have enduring value, reduce uncertainty and increase our understanding of when, where and how often the impacts of invasive species on biodiversity are indeed reversible.

## Introduction

Biodiversity is declining at global, regional and national scales (Dirzo et al. 2014; Diaz et al. 2019). Non-native invasive species are widely thought to be a major driver of this decline (e.g. Vitousek et al. 1997; Vilà et al. 2011; Doherty et al. 2016; but see Gurevitch and Padilla 2004). Beginning with Darwin’s visit in 1835, New Zealand has been considered a global exemplar of biodiversity decline caused by invasive species (Thompson 1922; Elton 1958; Allen and Lee 2006; Norton 2009; Simberloff 2019). Māori, as tangata whenua, have long expressed concerns about the plight of biodiversity and the undesirable role of invasive species (e.g. Lyver et al. 2008; Harmsworth and Awatere 2013). As a consequence, there is strong societal and political support for management to stop and reverse biodiversity decline. Safeguarding indigenous biodiversity is enshrined in national legislation (e.g. the Conservation Act 1987) and international obligations (e.g. the Convention on Biological Diversity 1993). This has led to some invasive species being managed for benefits to terrestrial biodiversity (Allen and Lee 2006; Jones and McGlinchy 2016; Hulme 2020). In recent efforts, the priority for expenditure has been mammal predator management and impact assessment (Peltzer et al. 2019; Hulme 2020). The Department of Conservation managed mammal predator and weeds on ca. 2.3 and 0.4 million ha respectively in the financial year ending June 2019 alone (Department of Conservation 2019). An underlying assumption for large-scale management is that biological invasions have caused declines in indigenous biodiversity and that controlling invaders will reverse these effects.

Benefits to biodiversity from invasive species management are widely anticipated in both policy (e.g. Parliamentary Commissioner for the Environment 2011; Predator Free 2020; Hackwell and Robinson 2021) and science reviews (e.g. Remeš et al. 2012; Byrom et al. 2016; Nelson et al. 2019; Binny et al. 2021; but see Caughley 1983; Hare et al. 2019). Moreover, these reviews, along with other sources of evidence, are required for decisions on invasive species management. The robustness of studies underpinning these evidential reviews has received little scrutiny (but see Simpkins et al. 2018), although some previous studies have noted deficiencies (e.g. Clayton and Cowan 2010; Smith et al. 2017) and recommended that greater robustness of evidence is desirable (e.g. Ferraro and Pattanayak 2006; Reddiex and Forsyth 2006; Doherty and Ritchie 2017; O’Grady 2020). More generally, the reliability of scientific evidence published in peer-reviewed journals requires ongoing scrutiny in several disciplines (e.g. medicine, Ioannidis 2005; biology, Baker 2016; social science, Camerer et al. 2018). New Zealand is rapidly scaling up invasive species management, as exemplified by Predator Free 2050 and the National Wilding Conifer Control Programme, and hence it is timely to assess the robustness of evidence used to infer biodiversity benefits from invasive species management.

Research evidence can be characterised as (in, arguably, increasing order of strength): anecdotes and casual observation; logical argument and mathematical modelling; systematic monitoring; and manipulative experiments (McArdle 1996; Hurlbert 1984). Manipulative experiments implemented as randomised control designs in human medical research have often been encouraged as the “gold standard” for causal inference (Jones and Podolsky 2015). In such experiments, ideally all variables are controlled for (accounted for by non-treatment measurements) and none are uncontrolled (McArdle 1996). If non-treatments work as predicted, it is possible to infer that the results are due to the effect of the treatment variable alone (Oksanen 2001; Ioannidis 2005); in our case the treatment of interest is an invasive species management action. Robust designs are necessary for resolving when and where the impacts of invasive species are, or are not, reversed by managing invasive species (see Coomes et al. 2003; Norton 2009; Doherty and Ritchie 2017).

In this paper we consider the robustness of field studies determining biodiversity responses to invasive species management in New Zealand. We first evaluate how key experimental design principles (hereafter “principles”; Table 1) have been implemented in recent studies testing the response of indigenous, terrestrial biodiversity (hereafter “biodiversity”) to manipulation of non-native invasive species (hereafter “invasive species”). Biodiversity responses included compositional, structural or functional characteristics of terrestrial ecosystems. Some principles underpinning robust designs are widely defined, taught and understood, but it is unclear how often these are applied. The manipulations are usually management actions (hereafter “treatment”). We evaluate peer-reviewed journal publications (hereafter “publications”), because journal publications are based upon long established scientific principles around repeatability and rigour. Studies published in the last decade (2010–2019) were identified by systematically searching selected journals, review articles, and using an internet search engine. Each publication was assigned one of three ordinal scores with respect to each of nine design principles (Table 1). The frequency of these scores, across all 155 publications, were used to interpret the robustness of recent field studies. We also determined whether the frequency distribution of scores (hereafter “score distribution”) for each principle were influenced by broad categories of invasive species or biodiversity responses. One expectation was that invasive species categories receiving high scores for principles would be those allocated relatively large expenditure (e.g., mammal predator). An alternative view is that high scores would be found for those organisms that are less challenging to sample in the field (e.g., stationary plants when compared to mobile birds). Finally, we briefly use these results to outline some pathways for improving the design of studies investigating biodiversity responses to invasive species management.

**Table 1.**
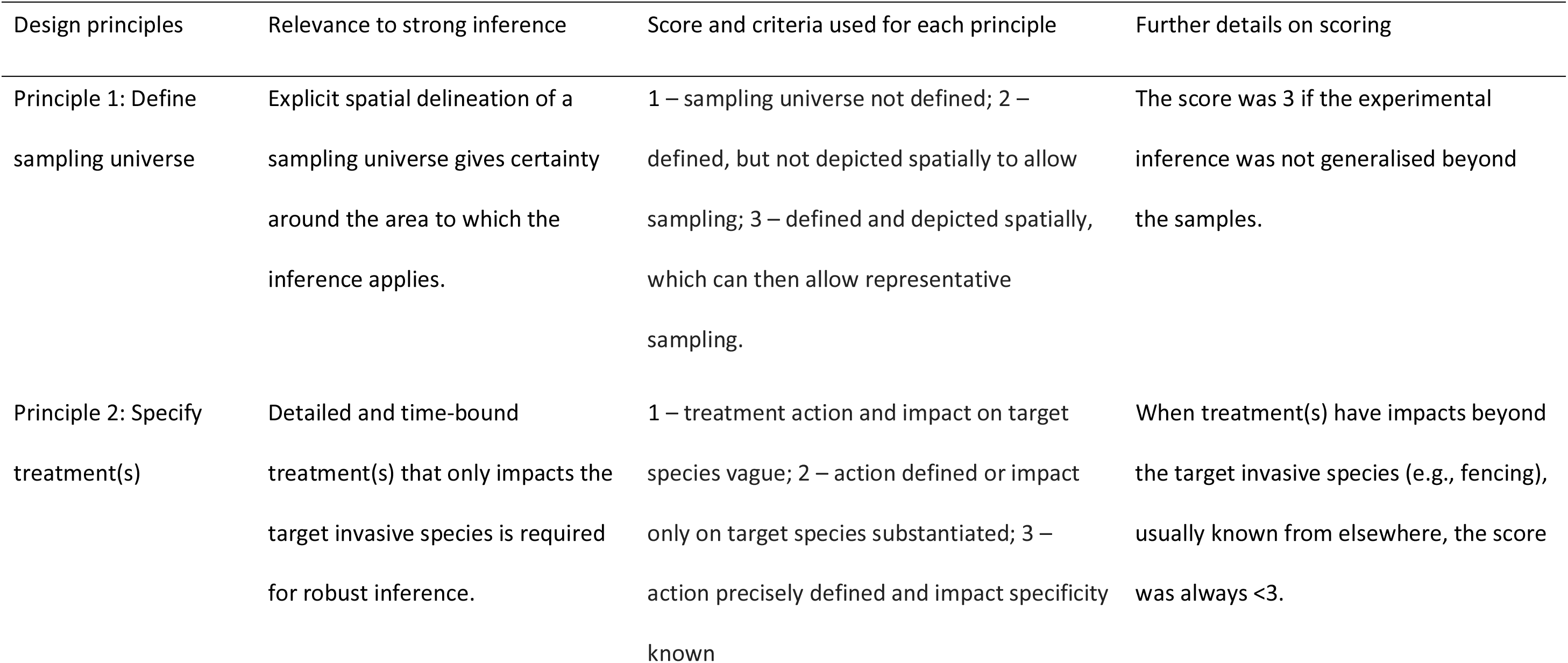
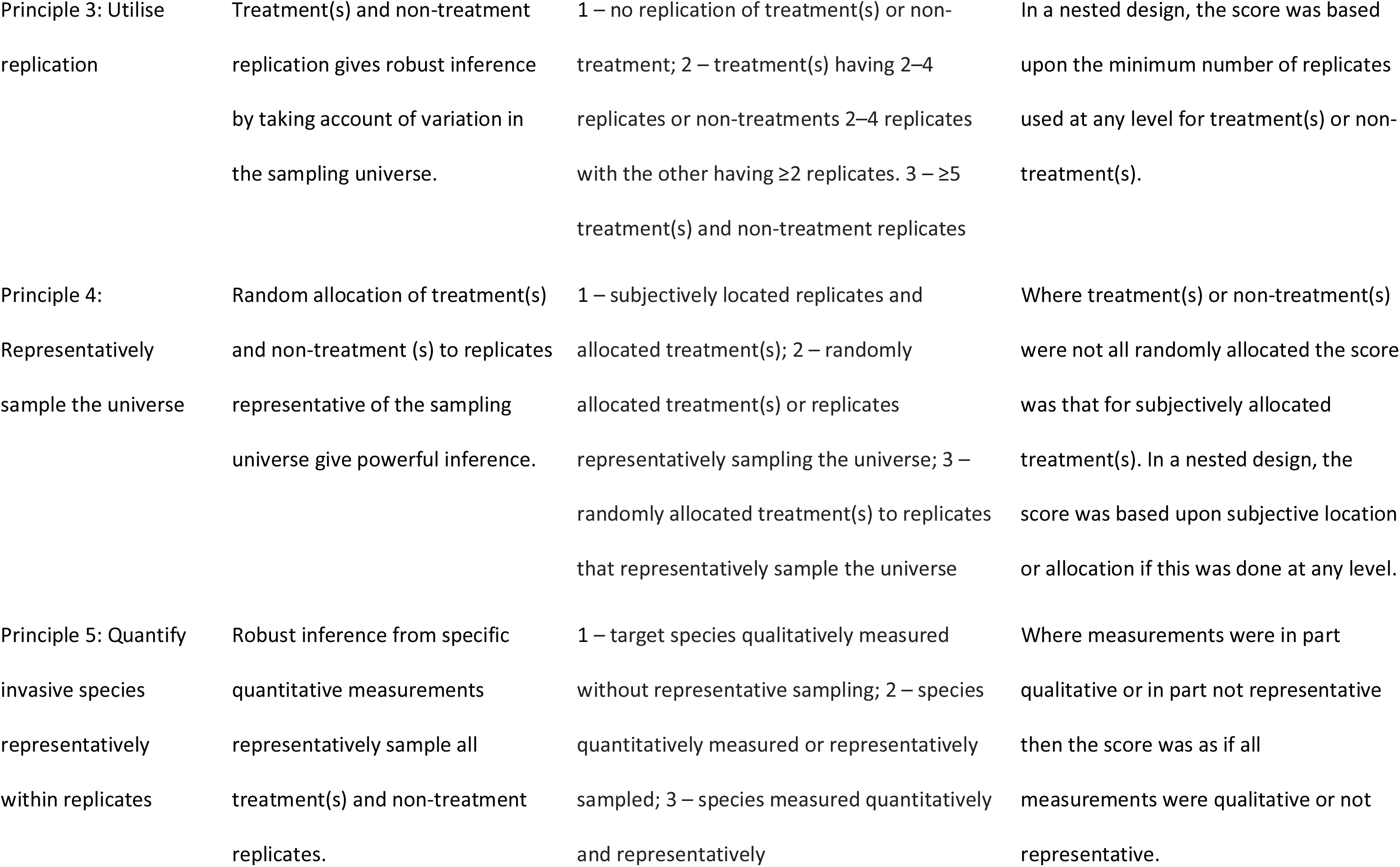
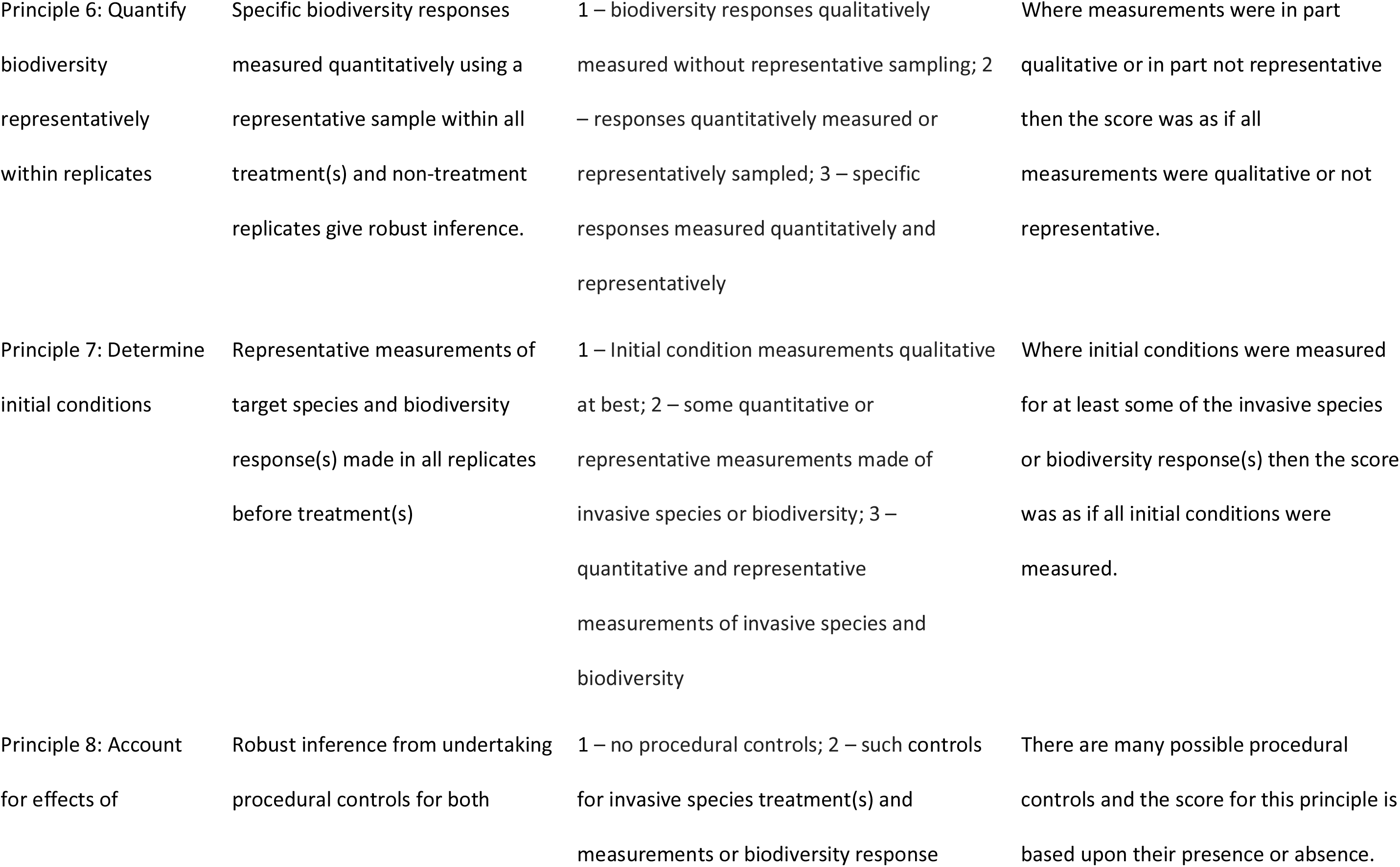
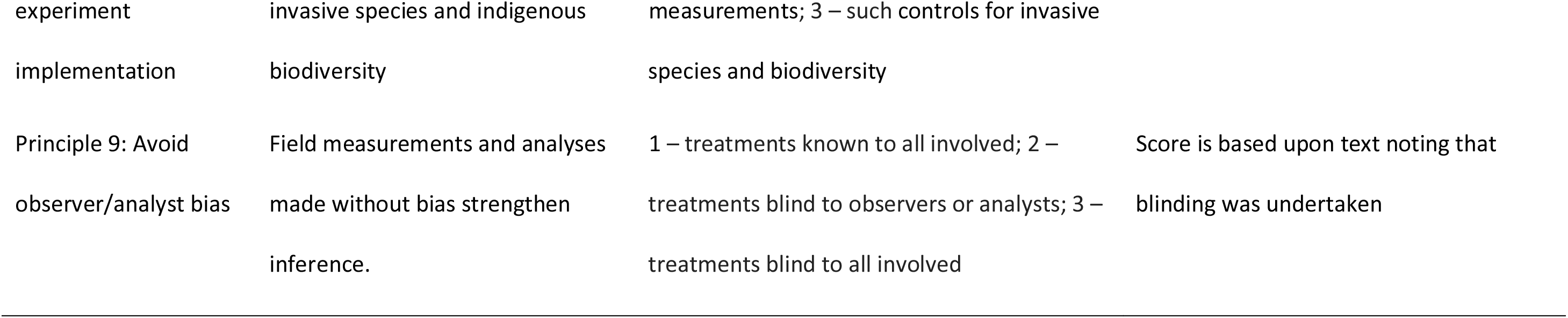
Experimental design principles used to evaluate each of 155 peer-reviewed journal publications. Also given are the relevance of each of the nine principles to the robustness of experimental inference, score and criteria used to evaluate each principle (high scores imply robust inference and vice versa) and further details on scoring.

## Methods used to assess robustness

### Selection of recent field studies

We selected publications that adopted a field experiment approach; that is, any field studies that derived a statistical inference from measurements of a biodiversity response to a treatment of one or more invasive species. Not all publications dealt with just one such experiment. Where part of a publication was based upon a field experiment, only that part was evaluated. Inferences were sometimes derived from multiple experiments within a single publication (e.g. experiments at multiple locations); in these few instances, all the experiments in that publication were evaluated using one set of scores. Although *in situ* biodiversity responses were commonly used, we also included experiments in which biodiversity was added (e.g. re-introductions). Types of treatments included any manipulation that attempted to directly affect invasive species at a particular location for a period of time. The majority of treatments were intended to reduce the abundance of invasive species or extirpate them from a defined area. We included studies in which invasive species were added. In some instances, for example fenced sanctuaries, the treatment occurred before biodiversity responses were measured. We included comparisons of islands with and without invasive species, as the absence of invasive species most likely reflects actions previously undertaken to restrict the movement of invasive species.

Our attention focussed on a field experiment approach because it forms a strong basis for inductive reasoning (Deaton and Cartwright 2018). In the first instance, we formed a list of 19 peer-reviewed journals where the authors’ considered such experimental work would be published (Appendix S1). Each publication, in each of those journals, was examined for relevance using issues published in 2010-2019. From this we selected 99 publications. These were then augmented by an additional 27 (giving a cumulative total of 126) publications, over the same years, found in the reference lists of 20 recent, relevant, review articles (listed in Appendix S2) identified by the authors. We then used various invasive taxa, biodiversity response and types of treatment, along with New Zealand, as search terms in Google Scholar at 30 April 2020 to select further publications, The search terms used were: weeds, fungi, bacteria, *Phytophora*, invasive species, mammals, and predators for invasive species; native plants, native invertebrates, fungi, bacteria, native bacteria, native fungi, native species, and biodiversity for biodiversity responses; and herbicide, biocontrol, control, fungicide, and antibacterial as types of treatment. These were used in various combinations (given in Appendix S2) and were broad terms to overcome the many possibilities (e.g., number of plant and bird taxa), but also to identify publications of less studied taxa (e.g., bacteria and fungi). From the Google Scholar search we selected a further 7 (total of 133) publications. Finally, from reference lists of all 133 publications we identified another 22 publications to evaluate (for a total of 155 evaluated (Appendix S1). The compiled publication list thus includes the vast majority of New Zealand field-based studies of invasive species management and biodiversity responses over the past decade.

### Scope of selected publications

The 155 publications were found in 46 peer-reviewed journals, with *New Zealand Journal of Ecology* the most common journal (23% of publications; Appendix S1). Mammals as predators were the most commonly studied invasive species category (58%) and native birds were the most commonly studied biodiversity response (43%). The most commonly studied relationships were between mammals as predators and birds (39%), and between mammals as herbivores and plants (19%). The first of these reflects a New Zealand priority of restoring threatened bird species (Hulme 2020). No study reported on potential relationships between mammal herbivores and bird responses despite these relationships being important elsewhere (e.g., Cocquelet et al. 2019; Crystal-Ornelas et al. 2021) and only two studies reported on relationships between invasive weeds and birds (Table 2).

**Table 2.**
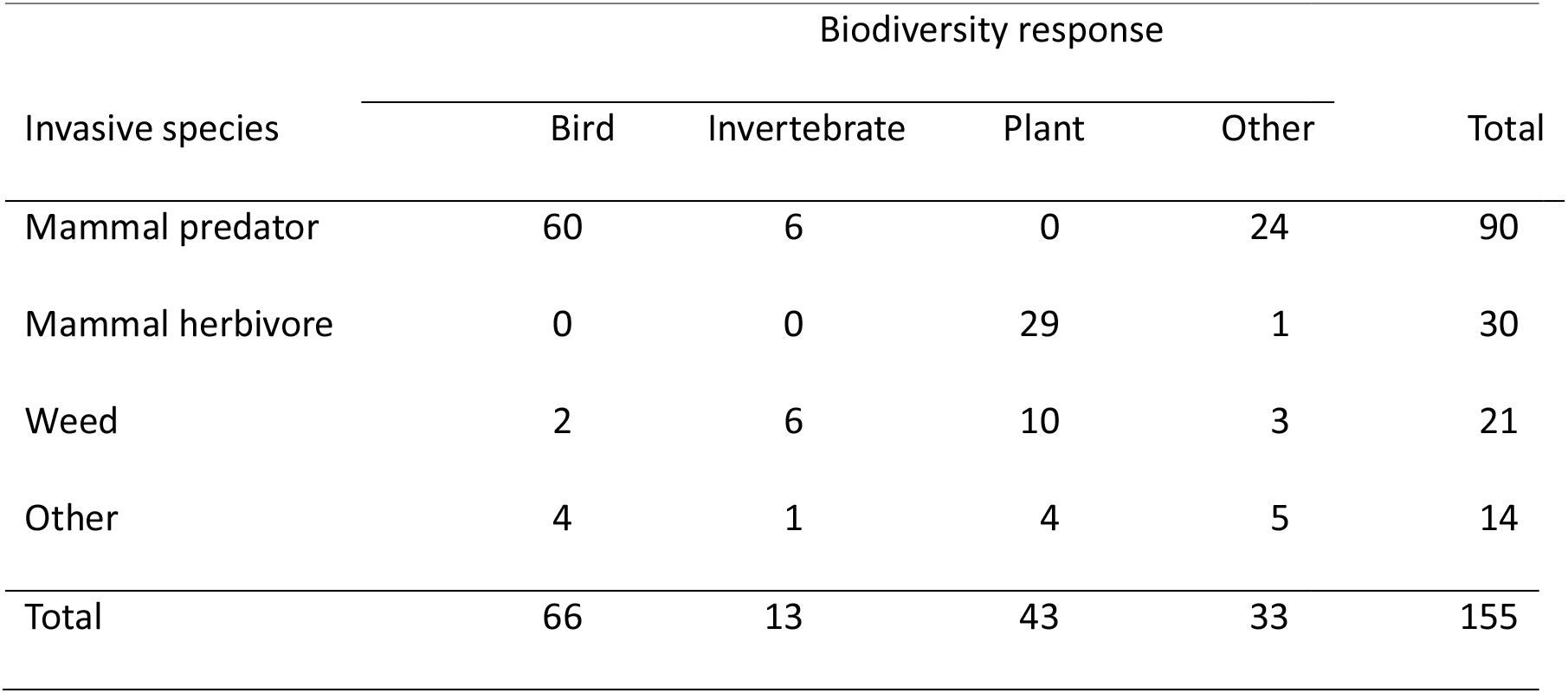
Number of peer-reviewed journal publications classified by invasive species and biodiversity response categories.

| Invasive species | Biodiversity response |  |  |  | Total |
| --- | --- | --- | --- | --- | --- |
|  | Bird | Invertebrate | Plant | Other |  |
| Mammal predator | 60 | 6 | 0 | 24 | 90 |
| Mammal herbivore | 0 | 0 | 29 | 1 | 30 |
| Weed | 2 | 6 | 10 | 3 | 21 |
| Other | 4 | 1 | 4 | 5 | 14 |
| Total | 66 | 13 | 43 | 33 | 155 |

#### Choice of principles

Each publication was evaluated against nine experimental design principles selected because of their contribution to the robustness of inference (Table 1; see also Scheiner and Gurevitch 2001). This list of principles was developed using two approaches. We first surveyed ecological syntheses and reviews that appraised experimental designs, but found these only focussed on some known principles (e.g., McArdle 1996; Hairston 1989; Hone 2007; Dickie et al. 2018). We then surveyed a wider literature to identify additional, relevant principles (e.g., Schulz et al. 2002; Ioannidis 2005; Greenlees et al. 2006). The nine emergent principles embody the majority of those pertinent to the design of ecological experiments. Furthermore, considering a greater number of principles than has typically been included in the evaluation of experimental design avoids potential bias from any one or few principles and broadens the range of principles considered in ecological research.

### Analyses

To determine the robustness of recent field studies, scores for all nine principles were assigned to each publication (Appendix S1), by one author (RBA), following an initial calibration among authors. We did this to ensure consistency of scoring (e.g. Baker 2016). Scores were made using an ordinal scale from 1 to 3 for each design principle, determined using the criteria and details in Table 1. The scores given for a principle were the lowest value justified by the publication. For example, it was assumed that treatments were not randomly allocated unless the article stated otherwise. The score distribution, for each principle, was determined across all 155 publications, where a high score (i.e. 3) implies more robust inference (Tables 1 and 3). Being across all publications means our score distribution for each principle represent the breadth of evidence.

**Table 3.**
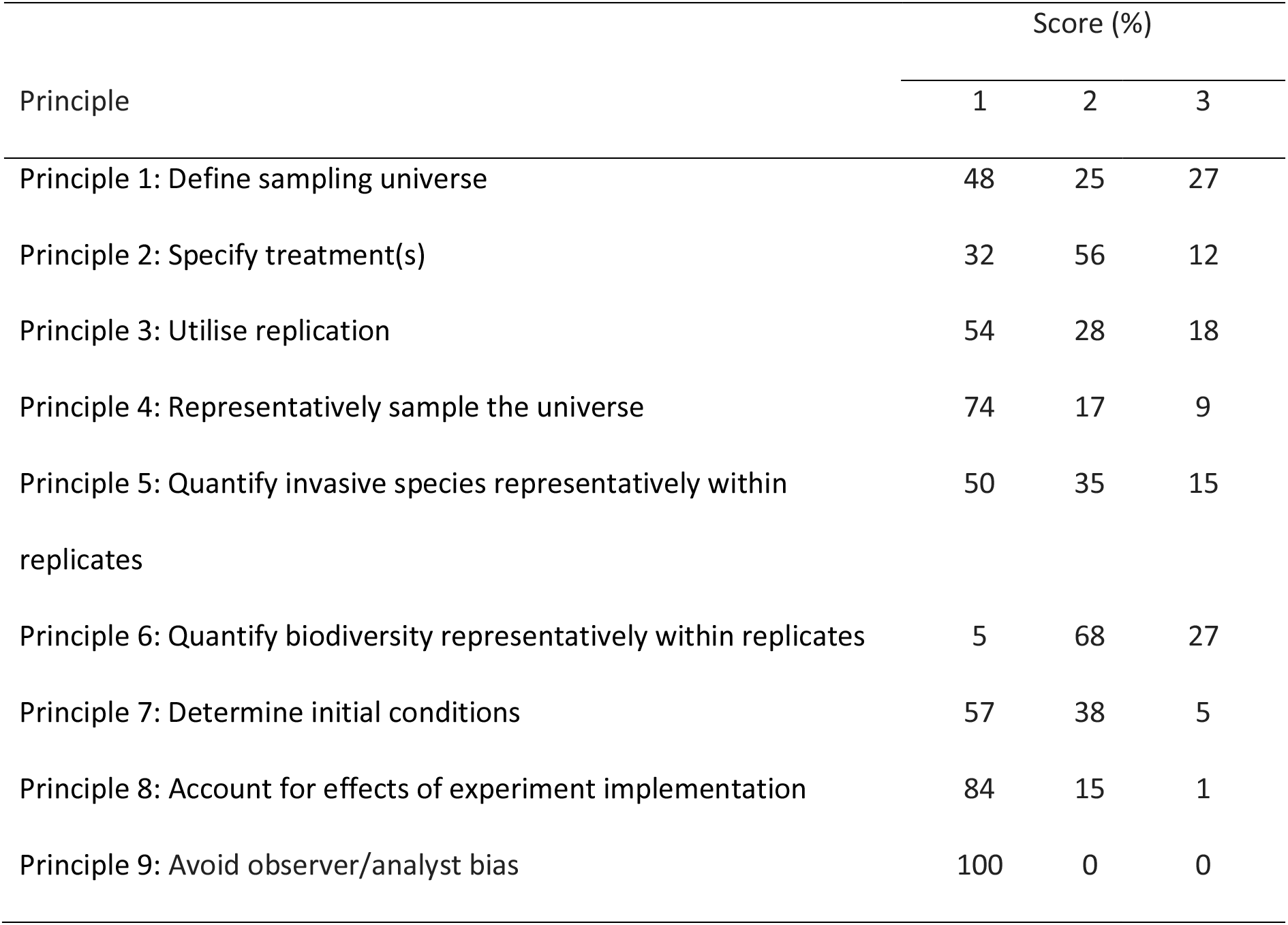
Percentage of 155 peer-reviewed journal publications receiving each score for nine experimental design principles. A higher score implies more robust inference.

We then assessed whether the score distributions were influenced by invasive species or biodiversity responses as covariates. Each covariate had four categories. Mammal predator, mammal herbivore, and vascular plant weeds (hereafter weed) were three categories used for invasive species, with bird, invertebrate, and vascular plant (hereafter plant) as three categories for biodiversity responses. Invasive species and biodiversity response covariates also included a fourth category, ‘other’. This category was a combination of two or three of the previously described categories, not one of the previously described categories, or a combination of previously described categories with one(s) not previously described. The other category for biodiversity responses included non-taxa related characteristics of ecosystems (e.g., soil properties). The score distributions for each of these four categories were based upon ≥13 publications and thus also represent a breadth of evidence (Table 2). As this breadth of evidence was not available for rarely studied invasive species or biodiversity responses our results are biased away from recently established invasive species. An invasive mammal species was included as herbivore mammal if a publication focussed on plant responses but as a predator if about animal responses. These categories defined invasive species and biodiversity response covariates which could explain variation in the score distribution for each principle. When presenting the results from the analyses of these covariates, one category is regarded as a reference category, and effect sizes for the other categories are estimated as the relative difference between that category and the reference category (on an appropriate scale; see below). Mammal predator and birds were regarded as the reference categories for the invasive species and biodiversity response covariates respectively, as they had the largest numbers of published studies.

Ordinal logistic regression was used to assess the effects of the two covariates on the score distributions for principles 1–6, and regular logistic regression for principles 7–8. Ordinal logistic regression can be used to assess the effects of covariates on the three scores, as there is a natural order to scores (Hosmer et al. 2013). The effects of the covariates can be interpreted in terms of odds ratios, in the same manner as for logistic regression. Regular logistic regression was used for principles 7 and 8, by combining the frequencies of scores 2 and 3, so that the score distributions had two values (i.e., score 1 and score 2/3 combined). The two scores were combined as there were very few publications with a score of 3 for principles 7 and 8. The data for principle 9 was not analysed further as all publications were assigned a score of 1 (Table 3).

Three models were fit to the data for each of the principles 1–8: no covariate effects; invasive species effects; and, biodiversity response effects. Models with both invasive species and biodiversity response effects were not considered as the covariates are not orthogonal (i.e., “Mammal predator” invasive species ware primarily studied in relation to “Bird” and “Other” biodiversity responses, while “Mammal herbivore” were almost exclusively studied in relation to “Plant” biodiversity responses (Table 2). Akaike Information Criterion (AIC) is an estimator of prediction error and, given a collection of models, enables the best model(s) to be identified (Burnham and Anderson 2002). Models were ranked on the basis of AIC to evaluate the level of support for each model and the relative importance of each covariate to each principle (Table 4). Models with a small difference in AIC (ΔAIC; relative to the top-ranked model) have a similar level of support to the top-ranked model, while models with larger ΔAIC values have much less support. A ‘small’ difference would be in the range of 0–2 AIC units, and a ‘large’ difference >4 AIC units. ΔAIC for the top-ranked model will always equal 0. When using logistic regression methods, estimated effect sizes may be interpreted as odds ratios (i.e., the multiplicative effect on the odds of a publication being given a particular ordinal score). The odds ratio was calculated from the logistic regression coefficient beta where the odds ratio equals exp(beta). Where beta equals 0 the odds ratio equals 1. An odds ratio equal to 1 is interpreted as no effect. For a covariate category (e.g. weed), an odds ratio not equal to 1 gives the relative number of publications that have a higher score, compared to the lowest score, than that shown by the relative number of publications in the reference category (e.g., mammalian predator for invasive species). If the odds ratio is >1 then more publications have a higher score and if the odds ratio is <1 then less publications have a higher score. The odds ratios, and overlap in associated 95% confidence intervals (CI), generated by the two forms of logistic regression were used to compare the effect of invasive species or biodiversity response categories on score distributions for each principle.

**Table 4.** Relative difference in Akaike Information Criterion value (ΔAIC) for each model compared to the highest-ranked model for the eight principles analysed by the two forms of logistic regression.

| Principle | Model |  |  |
| --- | --- | --- | --- |
|  | No covariates | Invasive species effects only | Biodiversity response effects only |
| Principle 1: Define sampling universe | 3.94 | 2.42 | 0.00 |
| Principle 2: Specify treatment(s) | 5.19 | 0.00 | 6.29 |
| Principle 3: Utilise replication | 44.70 | 0.00 | 17.69 |
| Principle 4: Representatively sample the universe | 0.00 | 1.05 | 4.42 |
| Principle 5: Quantify invasive species representatively within replicates | 9.74 | 0.00 | 9.33 |
| Principle 6: Quantify biodiversity representatively within replicates | 31.33 | 0.00 | 11.46 |
| Principle 7: Determine initial conditions | 0.33 | 0.00 | 2.50 |
| Principle 8: Account for effects of experiment implementation | 7.58 | 0.00 | 2.95 |

## Robustness of recent field studies

One model had more support than the other models considered for each principles, except for principles 4 and 7 (Table 4). For these principles, two models have similar levels of support. The model with the biodiversity response covariate most supported was for principle 1, and the model with the invasive species covariate most supported was for principles 2, 3, 5, 6, and 8. The model with no covariate effects was highest ranked for principle 4, although the model with the invasive species covariate also had some support. For principle 7, the invasive species effects model was ranked highest, but the model with no covariates had a very similar level of support. We now consider the most supported model within a wider consideration of the rationale for and evaluation of results for each principle.

### Define sampling universe

Principle 1 provides objectivity to the area to which the inference applies (Table 1; McArdle 1996). This universe can be expressed spatially at a point in time (e.g. locality, species distribution, or island) and, if carefully mapped, gives spatial limits. Forty eight percent of publications had the lowest score as they did not define the sampling universe and gave little detail on the area sampled (Table 3). The plant category scored lowest of the biodiversity responses, while the bird category tended to have relatively high scores (Fig. 1). The high score for birds possibly reflects publications where the birds were studied on easily defined areas such as an island(s) (e.g. Jones et al. 2015) or an area fenced to exclude predators (e.g. Bogisch et al. 2016). The utility of publications which do not define the sampling universe is limited, because their results cannot be interpreted in relation to an area over which they apply or extrapolated to larger areas (Smith et al. 2017). Such a limitation appears common in ecological research. Dickie et al. (2018), for example, found that 92% of 75 DNA-based biodiversity studies from around the world did not define a sampling universe; often studies described sampling locations in detail, but not how these locations were chosen to be representative of any larger area.

**Figure 1.**
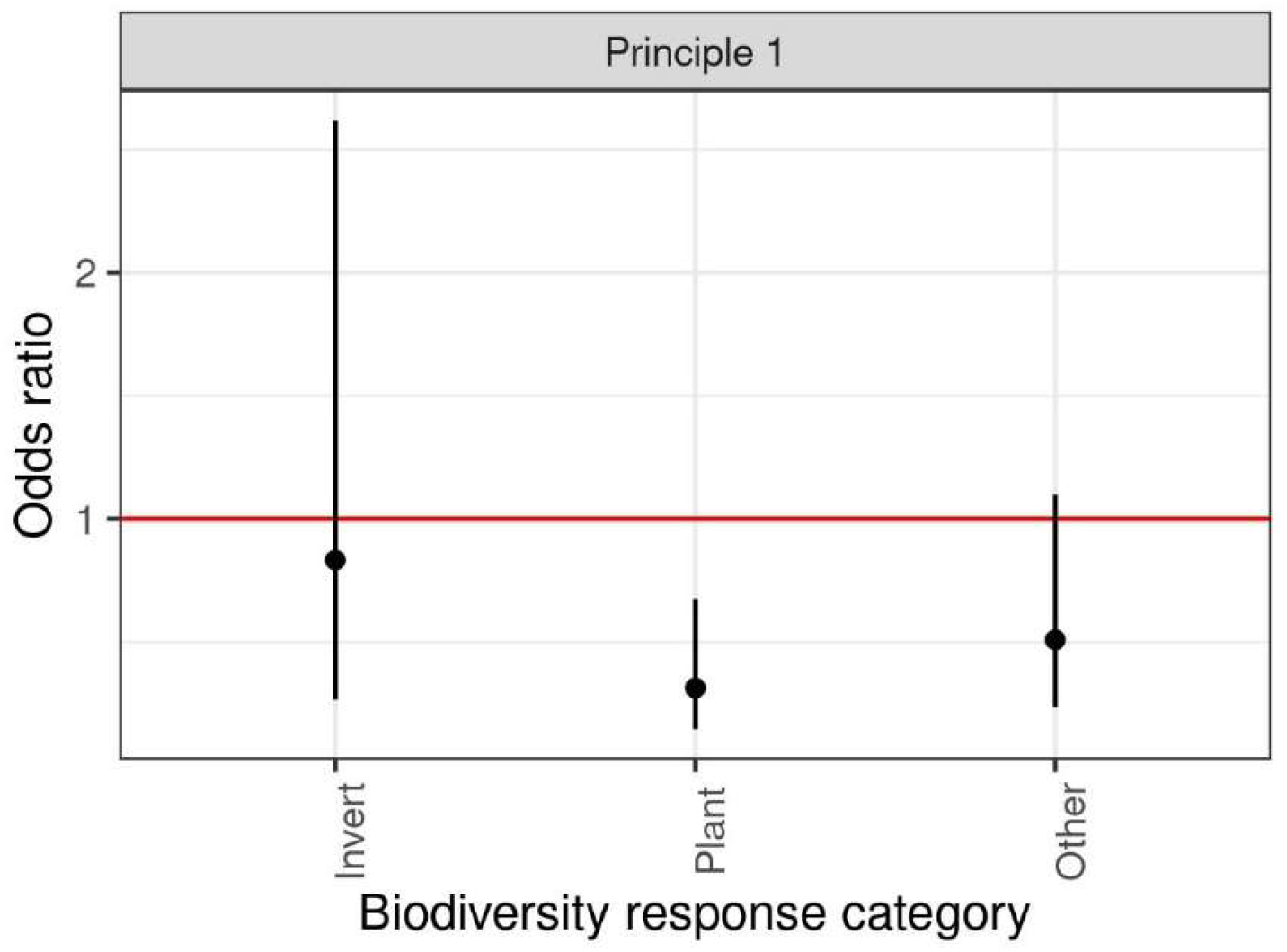
Estimated odds ratios (and 95% confidence intervals) for biodiversity response categories. The best supported model for Principle 1: Define sampling universe was for the biodiversity response covariate (Tables 1, 2 and 4). The reference category was birds (horizontal line). Invert, Invertebrate.

### Specify treatment(s)

A time-bounded treatment should be defined with sufficient methodological detail to provide clarity about what is being tested; moreover, the treatment should only impact the target invasive species and not be confounded with other actions (Principle 2). Fifty-six percent of publications received an intermediate score for this principle, because non-target invasive species or biodiversity itself were potentially influenced by the treatment(s), rather than the treatment action not being defined (Tables 1 and 3). The use of non-target specific poisons as treatment(s) for mammal predator and mammal herbivore may explain why they have lower scores than weed or other invasive species categories (Fig. 2). Moreover, field experiments were at times confounded by the addition of biodiversity to some invasive species treatments or replicates: these included additions of native species of birds (e.g. Bombaci et al. 2018), plants (e.g. Graham et al. 2013) and reptiles (e.g. Tanentzap and Lloyd 2017) to treatments. In such cases, any biodiversity response, in certain treatment(s) or replicates, may be directly, or indirectly, a consequence of the addition(s). Some experiments were also confounded by the treatment(s) being applied previously, at least in part, to non-treatment(s) (Fea and Hartley 2018), or the treatment(s) changing during the course of a study. Invasive mammal species treatment(s), for example, should not change trapping and poisoning regimes during a study, because they differ in both effectiveness of control and non-target impacts (e.g. Parkes and Murphy 2003). There is the potential for hidden treatments to create complex interactions that drive ecosystem-level responses (e.g. Wardle et al. 2012). As a consequence, scores for this principle were high where a manipulation added only the target invasive species (e.g. Pawson et al. 2010) or affected only the target species (e.g. Morgan et al. 2012).

**Figure 2.**
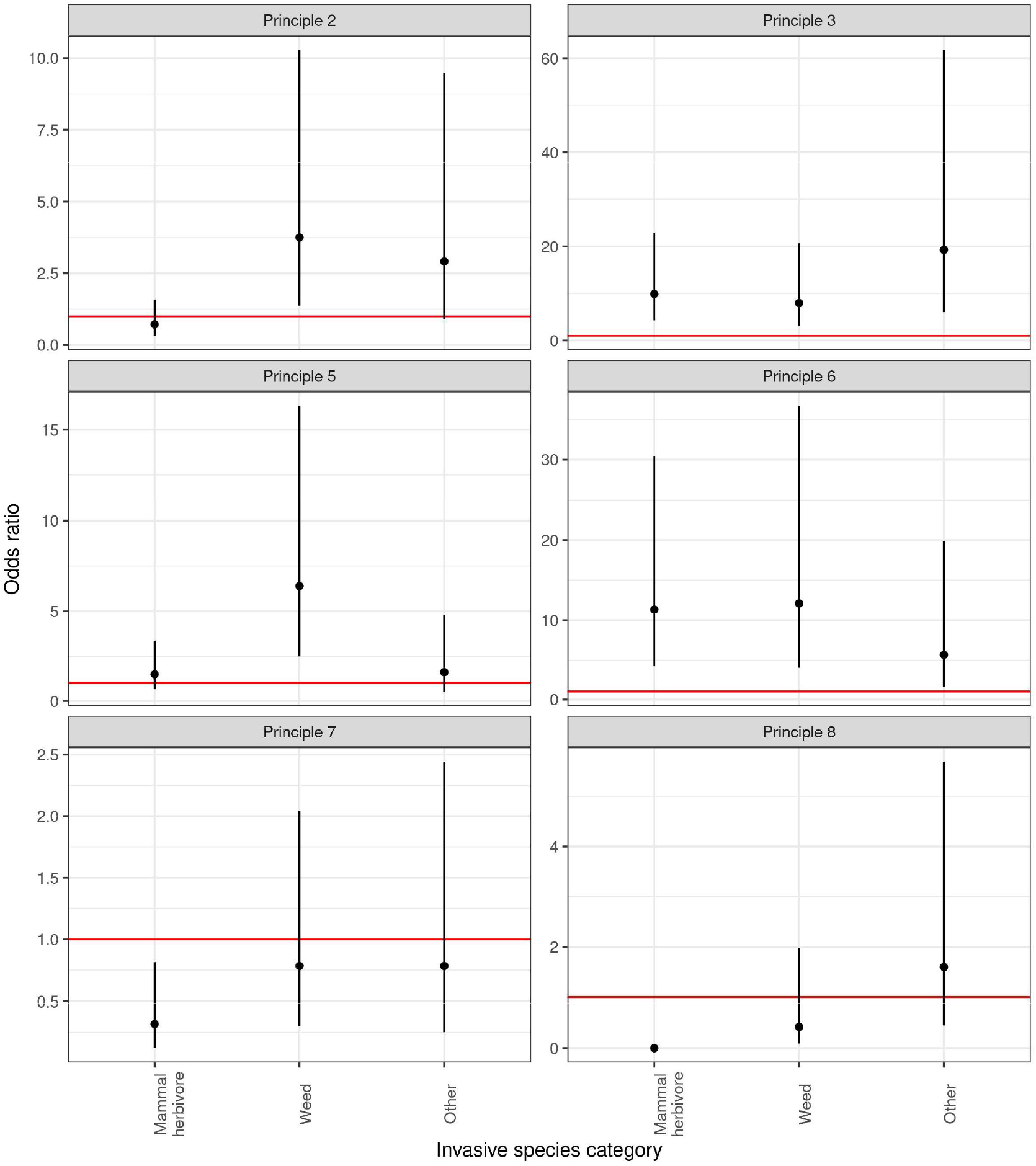
Estimated odds ratios (and 95% confidence intervals) for invasive species categories. The best supported models for Principles 2: Specify treatments, 3: Utilise replication, 5: Quantify invasive species representatively within replicates, 6: Quantify biodiversity responses representatively within replicates, 7: Determine initial conditions, and 8: Account for the effects of experiment implementation was the invasive species covariate (Tables 1, 2 and 4). The reference category was mammal predator (horizontal line).

### Utilise replication

Treatment and non-treatment replicates are required in order to adequately account for variation found within a sampling universe (Principle 3; Hurlbert 1984; Chalcraft 2019). Replication is challenging for some field experiments, but it is often useful to have more than four replicates to reliably detect treatment effects (Carpenter 1989; Eberhardt and Thomas 1991). Fifty-four percent of studies evaluated had no replication of treatment(s) or non-treatment(s), and only 18% had enough replicates to reliably detect treatment effects (Table 3). Twelve percent of publications considered one treatment location, such as an island or fenced sanctuary, and without non-treatment locations. This may be why mammal predator studies, often treated as such locations, had relatively low scores (Fig. 2). While it is particularly challenging to replicate mammal herbivore studies they were replicated as well as weeds (Fig. 2). Relatively few replicates could be required if there is little variation in treatment(s) and non-treatment(s) (Oksanen 2001). The level of replication required can be clarified by power analyses (Allen et al. 2003). However, New Zealand is remarkably diverse in climate, soils, geology, and disturbance history, even at small spatial scales (e.g. Wardle 1991). For example, fresh leaf phosphorus (P) concentrations subjectively sampled in Te Urewera, central North Island, captured >90% of the global variation in leaf concentrations (Richardson et al. 2008). In Te Urewera, this variation reflected topographically related differences in soils at the scale of hundreds of meters. Not only does this small-scale variation drive marked differences in nutrient cycling (Richardson et al. 2004), but also habitat use by invasive species such as feral pigs (Forsyth et al. 2016). Elsewhere, Forsyth et al. (2015) have suggested, through a combination of empirical studies and modelling, that the long-term impacts of invasive deer and rodents on indigenous forests will be greater on sites with relatively high soil P availability than nearby sites with low P availability. Hence, robust replication and randomisation are needed (Hurlbert 1984; Filazzola and Cahill 2021). Chance intrusions also occurred in some replicates during an experiment, which added variability to the data. Such intrusions can include flood damage (e.g. Simpkins et al. 2015), predation by domestic dogs (e.g. Robertson et al. 2019), build-ups in disease causing fungi (e.g. Perrott and Armstrong 2011), and variation in avian malaria infection rates (e.g. Alley et al. 2010).

### Representatively sample the universe

It is desirable that both treatment and non-treatment replicates, which can occur at multiple levels in a complex design, are representative of the sampling universe, either randomly or systematically located, and have randomly allocated treatment(s) (Principle 4; Mentges et al. 2021). Seventy-four percent of studies neither had replicates that representatively sampled a sampling universe, nor employed a random allocation of treatment(s) (Fig. 3; Table 3). Twenty percent of publications randomly allocated treatment(s) to replicates. It is also desirable to have treatment(s) and non-treatment(s) replicates intermixed within the sampling universe to accommodate any spatial patterns (Hurlbert 1984). This requires replicates to be independent, i.e. sufficiently distant, to avoid spatial autocorrelation or spill-over of treatment effects. Some publications evaluated acknowledged interchange between treatment and non-treatment replicates (e.g. Parlato and Armstrong 2013).

**Figure 3.**
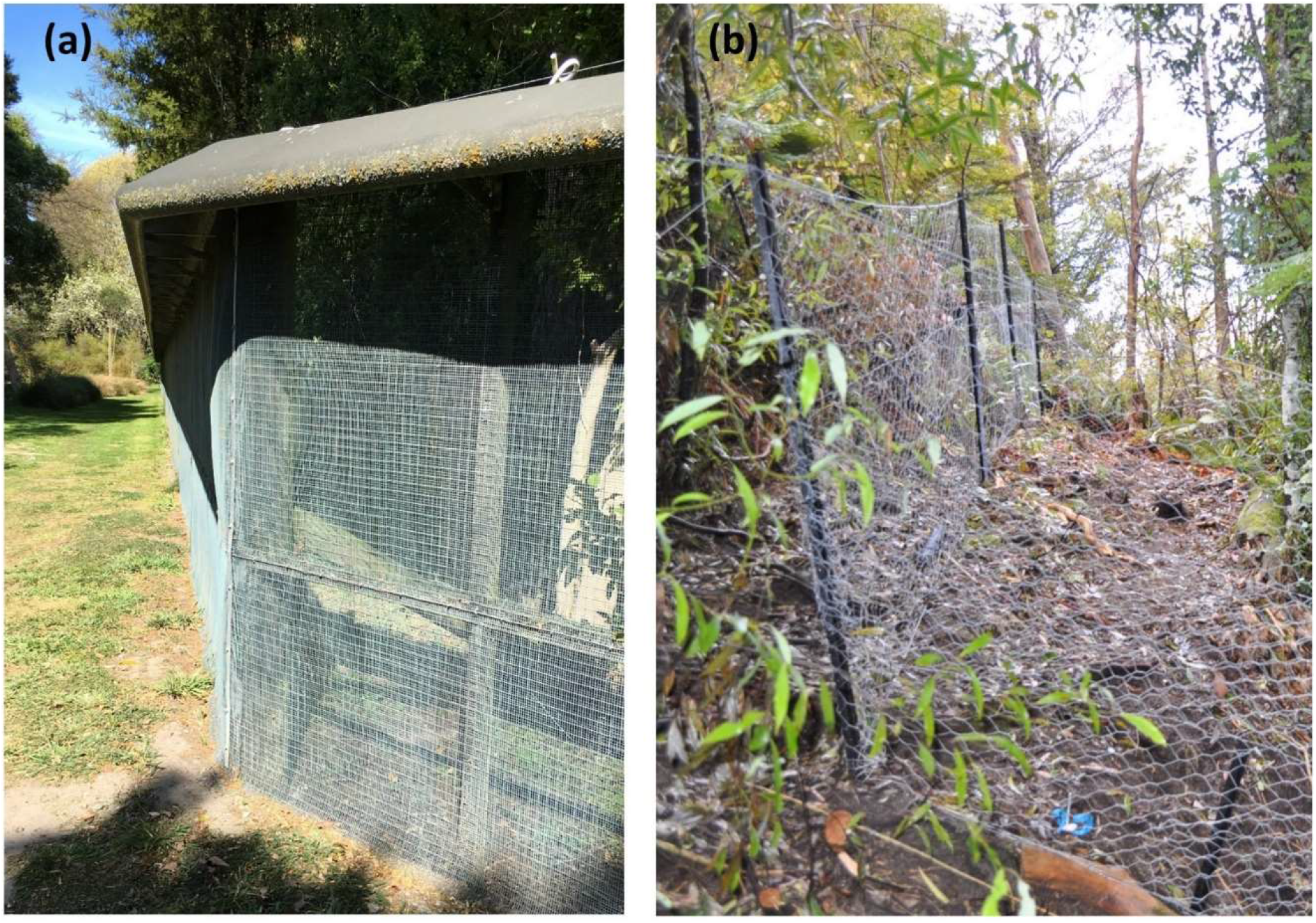
A fence constructed with a fine wire mesh, capped, and topped with an electric wire has been used at Riccarton Bush, Christchurch, to exclude invasive mammal predators such as brushtail possums, stoats and rats (a). Simpler fences constructed with a coarse wire mesh have been used in Te Urewera to exclude invasive mammal herbivores such as deer (b). Such fencing treatments are widely deployed for animal exclusion but often exclude non-target species, have been located subjectively, treat a small area, are subject to periodic breaches, and potentially influence ecosystem-level responses (e.g. seed production and litter fall).

Management imperatives or financial and logistical constraints have sometimes driven suboptimal treatment(s) allocation (e.g. Parkes et al. 2006; Broadbent et al. 2017). Such subjectivity accentuates potential problems of hidden treatment effects and sampling bias (Smith et al. 2017). If this bias is systematic, then they become embedded in meta-analyses and reviews. In a rare example, Peltzer et al. (2014) shed some light on an apparent systematic bias in fenced exclosure studies that explore biodiversity responses to removing invasive ungulate herbivore species (Fig. 3). Conservation managers subjectively established ca. 100 fenced exclosures throughout New Zealand’s indigenous forests during the 1970s and 1980s to remove ungulate browsing in the forest understorey (Mason et al. 2010). As expected, palatable tree species showed higher numbers of small trees inside exclosures than outside (Mason et al. 2010). Unexpectedly, the numbers of these palatable species found on a representative grid of locations (unfenced) nationally were similar to those found inside the fenced exclosures (Peltzer et al. 2014). It appears that the exclosures were constructed, knowingly or unknowingly, on sites where ungulate impacts were strongest. Under such circumstances, any level of replication (number of fenced exclosures), for example, would be overridden by sampling bias.

### Quantify invasive species representatively within replicates

Representative sampling of replicates and quantitative measures of invasive species (Principle 5) ensures robust inferences (Oksanen 2001; Salo et al. 2010). In treatments where invasive species have previously been eradicated (e.g. islands), our evaluation of Principle 5 partly used support from existing literature cited, rather than publication-specific measurements. Fifty percent of publications reviewed neither representatively sampled nor quantitatively measured the target invasive species (Table 3). In total, 84% of publications did not use representative sampling (see implications in principle below), and 51% of publications did not include quantitative measurements of target invasive species. We had expected that weeds would score relatively high for principle 5 (Fig. 2), because such stationary organisms are relatively easy to measure. In some publications, quantitative measurement was not undertaken because the target invasive species was assumed to be eradicated or never present; this assumption needs to be verified, because ongoing (re)invasion occurs and can be difficult to detect (e.g. Bellingham et al. 2010; Drummond and Armstrong 2019). In other publications, the manipulation itself was assumed to cause an invasive species difference between treatment(s) and non-treatment(s) replicates. However, Forsyth et al. (2013) showed that there can be large uncertainty around the impacts of an invasive species manipulation in treatment and non-treatment replicates. Salo et al.’s (2010) review stressed the importance of measuring the abundance of predators before and during an experiment to be sure about their impact on prey.

### Quantify biodiversity responses representatively within replicates

Representative sampling of replicates and quantitative measures of biodiversity responses (Principle 6) also ensures that inferences are robust (Oksanen 2001; Salo et al. 2010). Sixty-eight percent of publications received an intermediate score for this principle (Table 3). This is because publications were often based upon quantitative measurements of biodiversity responses but not representative sampling. In quantifying biodiversity responses, mammal predator studies tended to score relatively low when compared with the scores for the other three invasive species categories (Fig. 2). Twenty-seven percent of publications described representative sampling (e.g. plots) within replicates (Table 3). Representative sampling remains poorly utilised, even in the face of recent technological developments that simplify its use (e.g. Geographic Information Systems; Smith et al. 2017). Rather, subjective sampling was the norm in our reviewed publications, and this has important implications. Avoiding certain locations or features within replicates may reduce variability, and likely increases the probability of significant effects (Dickie et al. 2018). For example, McAlpine et al. (2016) subjectively avoided invasive pine slash sites in treatments when measuring native seedling establishment. Such an approach means that results are only representative of a subset of the sampling universe, and this reduced sampling universe needs to be defined.

### Determine initial conditions

If few replicates used in an experiment, or background variation among replicates is large, then inferences can be strengthened by accounting for initial conditions. One way of doing this is by using before-and-after treatment(s) measurements of invasive species or biodiversity responses (Principle 7; Hurlbert 1984; Hairston 1989; Salo et al. 2010). In our evaluation, before treatment measurements were interpreted as representing initial conditions rather than non-treatments. Fifty-seven percent of publications did not measure initial conditions in field experiments (Table 3).

Mammal herbivore studies received a relatively low number of high scores (Fig. 2). Measuring initial conditions is particularly challenging and expensive for wide ranging mammal herbivores. Where initial conditions were measured, sometimes seasonal measurements were more restricted before a treatment than after (e.g. van Vianen et al. 2018), or for only a subset of biodiversity responses (e.g. Robertson et al. 2019). Representative and quantitative measurements of initial conditions were rarely made for both biodiversity responses and invasive species (Table 3). Indeed, manipulations of invasive species were at times undertaken long before any measurements, with long-term vegetation change creating a challenge for understanding any variability in initial conditions (e.g. Holdaway et al. 2014).

### Account for effects of experiment implementation

Procedural controls ensure that the observed effect is not an artefact of some aspect(s) of experiment implementation (Underwood 1997). These were evaluated for both target invasive species and biodiversity responses (Principle 8). Procedural controls were rarely used (84% scored 1; Table 3). Mammal herbivore publications scored relatively low for principle 8 when compared to the other three invasive species categories (Table 3; Fig.2). Non-target impacts of various poisons, used to manipulate invasive mammals, on biodiversity has forced some scrutiny of the unintended poisoning of native animals as a procedural control (e.g. Kemp et al. 2019). Another example examined whether bird translocations to islands without predators caused a directional selection on a stress response to capture (Adams et al. 2013). Publications with procedural controls tended to publish those results alone, without including the invasive species and biodiversity responses to treatment(s). Only the study by Wardle et al. (2010) included more than one of the many possible procedural controls. The importance of procedural controls has long been recognised in ecological research (Hurlbert 1984; Greenlees et al. 2006), yet is poorly developed when compared with its use in medical research (Price et al. 2008). There are many possible procedural controls and individual studies need to consider what effects to account for in experimental implementation.

### Avoid observer/analyst bias

We include treatments being blind to observers and analysts (Principle 9; Schulz et al. 2002) as our final principle. Implementing and reporting experiments that are blind is a key way of avoiding bias (Ioannidis 2005). All publications used field experiments in which the treatment(s) were apparently known to the observer and analyst (Table 3) which could introduce bias in data collection, analysis or interpretation. This allows for considerable bias when, for example, treatment effects on biodiversity are not quantified but subjectively estimated. The absence of observer/analyst blinding contrasts with its common use in other research areas as a means of overcoming bias (Schulz et al. 2002). There could also be a role here for independent audits of field measurements.

## Pathways for improvement

### Enhancing field studies

Experiments require substantive assumptions, prior information, and are not independent of “expert” knowledge (Deaton and Cartwright 2018). We support a greater commitment to robust experimental designs, but recognise that there is a trade-off between this and the greater logistical and financial ease of simple designs (Christie et al. 2019; Filazzola and Cahill 2021). As a consequence, our intent is to improve the robustness of future field studies for at least some principles. Crawley (2015) offers some guidance and considered randomisation and replication as the two essential concepts in experimental design. We discuss below examples of how well-known principles could be better implemented in field studies.

### Clearer protocols for some principles

Some design principles such as replication and randomisation are well established and could readily be applied more widely. For example, large-scale invasive species management could use random allocation of control treatments. Other principles such as procedural controls and minimising observer/analyst bias are more challenging to utilise in ecological research. Implementing these challenging principles requires a deconstruction of the experiment to understand what is required. Implementing an experimental manipulation could create, for example, artefacts that directly or indirectly affect both target invasive species, other invasive species, the biodiversity response of interest, or other biodiversity responses. Not all of these will necessarily be important for procedural controls, and this may change through time. Testing whether implementing a manipulation itself directly influences an invasive species may require non-treatment(s) to receive only the action used in a treatment, for example, shooting at an invasive ungulate species using blanks in the non-treatment areas in an experiment testing the effects of aerial shooting. Procedural controls are challenging in a common New Zealand manipulation, fencing, whereby there is a direct effect on the target invasive species as well as other invasive species. Fences were commonly used to exclude invasive mammal predators to restore bird populations, but they often also exclude invasive herbivores (Fig. 3). If the herbivore exclusion enabled forest understories to become denser, then reduced visibility may mean less bird nest predation and increased bird abundance (Cocquelet et al. 2019; Crystal-Ornelas et al. 2021). Understanding such relationships in New Zealand would benefit our interpretation of mammal predator/bird relationships. Procedural controls, and specifying the treatment, can be improved when the target invasive species is added as a treatment.

### Greater scrutiny pre-implementation

Some mechanisms already exist for greater scrutiny before a field experiment is implemented. Pre-registering a study is now becoming an accepted part of conservation science (Parker et al. 2019; Filazzola and Cahill 2021). Pre-registration is the archiving of a detailed description of a proposed study’s research questions/hypotheses, experimental design, and data collection and analysis methods in a public registry. Registered reports are similar to pre-registration, but are subject to peer review at a journal prior to the study beginning. If after peer review the journal editor is satisfied with the logic and design of the proposed study, then in-principal acceptance of the study is given, regardless of results, provided that the robust design and analysis described in the registered report were followed. Some ecological journals offer the registered report option (e.g. *Conservation Biology*). While such options may not yet be appropriate for all studies, it is likely that preregistration and registered reports will become an increasingly accepted way of doing ecological research and are one mechanism to help ensure that key design principles are addressed in field studies.

### Adequately resourced experiments

A lack of resources in ecological research and monitoring can lead to a weak and biased evidence base (e.g. Dirzo et al. 2014; Christie et al. 2019). This limits our ability to provide robust recommendations to decision-makers. Greater resources is not the only solution, and one alternative is to undertake fewer experiments that are individually more expensive because they better adhere to design principles (Baker 2016; Christie et al. 2019; Filazzola and Cahill 2021). For example, this may allow replication (principle 3) and quantification of biodiversity responses (principle 6) to be strengthened in mammal predator studies as well as initial conditions (principle 7) and procedural controls (principle 8) to be more commonly applied in mammal herbivore studies (Fig. 2). Fewer experiments could simply be generated by, for example, using registered reports, as defined above, to improve design prior to treatment or data collection.

### Strengthen publication processes

Reviewers, editors and journals could be more insistent that publications meet widely known and accepted design principles (Filazzola and Cahill 2021). Our evaluation suggests that design robustness may not be a key decisive factor when manuscripts are accepted for publication in a wide range of journals. We suggest that citations of individual publications be qualified by denoting their critical design principle limitations (e.g. “subjectively sampled” or “unreplicated”). When meta-analyses are undertaken, weighting systems can be used to give greater influence to individual studies with more robust designs (Stewart 2010; Christie et al. 2019). However, increasingly elaborate analyses cannot remove potentially widespread systematic bias (Peltzer et al. 2010). Increased transparency about robustness, uncertainties or bias could strengthen their inclusion in decision-making and, moreover, reinforce the utility of having robust experiments as an evidence base.

### Adopting other approaches

We support a view that questions a universal role for experiments as an inductive approach because of: “big-data” mining opportunities, the complementary role of deductive reasoning, traditional knowledge, and design challenges when implementing robust experiments (e.g. Eberhardt and Thomas 1991; Oksanen 2001; Lyver et al. 2008; Jones and Podolsky 2015; Deaton and Cartwright 2018; Munafò and Smith 2018; Kreyling et al. 2018; Filazzola and Cahill 2021). Circumstances where it is difficult to replicate an experimental manipulation reinforce the case for various forms of systematic monitoring (Hurlbert 1984; Eberhardt and Thomas 1991). However, most of the design principles outlined above (e.g. sampling universe, representativeness) also apply when generating evidence from such approaches. It is for such reasons, for example, that the Department of Conservation’s Biodiversity Monitoring and Reporting System samples New Zealand’s indigenous forests nationally on a grid with a random starting point (Allen et al. 2003; Forsyth et al. 2018; Bellingham et al. 2020). Such approaches can generate insights at larger spatial and temporal scales than experimental studies, thereby increasing the generalisability of results, but also have limited ability to disentangle different drivers of biodiversity responses. It is therefore desirable to include interpretive covariates in systematic monitoring studies (Allen et al. 2003), although experiments do not necessarily relieve us of this need (e.g. Deaton and Cartwright 2018). Interpretive covariates focus our attention away from typical effects shown in experimental studies onto research that accommodates idiosyncrasies and differences (e.g. Kreyling et al. 2018). Context matters for invasive species impacts (e.g. Sapsford et al. 2020). With such approaches, alternative analytical methods such as noise clustering of community composition (e.g. Wiser and de Cáceres 2013) and statistical matching (e.g. propensity scoring) can be used, for example, to investigate the effects of manipulating invasive species on biodiversity (Ramsey et al. 2019). These methods require some treatment(s) locations to have covariate values that overlap with non-treatment locations. This overlap is achieved for common invasive species and biodiversity when sampled at large spatial scales, as well as capturing spatial structure (Legendre et al. 2004).

## Concluding comment

As field ecologists, we acknowledge the difficulties of implementing and sustaining field studies at meaningful spatial and temporal scales, and studies that we have recently published were evaluated in this review (e.g. Peltzer et al. 2014; Ramsey et al. 2017) and scored low in some design principles (Appendix S1). Nevertheless, our review highlights both long-recognised (e.g. replication and randomisation; Hurlbert 1984) and more recent (e.g. blinding; Schulz et al. 2002) design principles not yet widely adopted in ecological field experiments. Other methodological areas not considered here, such as suitability of statistical analyses, incorporating measurement error, role of audit, utility of response variables, code sharing and data accessibility (Salo et al. 2010; Fanelli 2012; Tressoldi et al 2013; Fraser et al. 2018; Mason et al. 2018; Cusser et al. 2021; Filazzola and Cahill 2021), also underpin the utility and robustness of publications. Incorporating measurement error, for example, can alter conclusions drawn from biodiversity monitoring (Mason et al. 2018). However, we emphasise that robustness of design precedes or underpins many of the other methodological areas mentioned above. Research on research, as we have undertaken, is necessary (Ioannidis 2018). The importance of this issue cannot be overstated: ongoing declines, in some components of biodiversity, despite more than a century of management efforts, suggest that greater and robust knowledge are essential to inform management (e.g., Coomes et al. 2003; Peltzer et al. 2019; Simberloff 2019; Hulme 2020). Increasingly robust studies will allow us to better understand when, where, and how often the impacts of invasive species will persist or be influenced by management.

## Supporting information

Supplemental searches

## Acknowledgements

We thank Chris Jones and the Predator Free New Zealand Trust for providing access to an unpublished report. This article was funded by the authors and Department of Conservation. John Herbert, Sharyn Laing, Murray Llewellyn and Elaine Wright provided motivation for this review and Sarah Richardson the photograph of a fence excluding invasive ungulate herbivores. We thank three anonymous reviewers for comments that greatly improved this manuscript.

## Data and code availability

All data is made available in Appendix S1. Code is available through **Darryl I. MacKenzie**.

## Author contributions

**Robert B. Allen**: Conceptualisation, Methodology, Investigation, Original Draft. **David M. Forsyth**: Conceptualisation, Methodology, Review and Editing. **Darryl I. MacKenzie**: Conceptualisation; Analysis and Review. **Duane A. Peltzer**: Conceptualisation, Review and Editing, Visualisation.

